# Identification of an allele-specific transcription factor binding interaction that regulates *PLA2G2A* gene expression

**DOI:** 10.1101/2023.12.12.571290

**Authors:** Aki Hara, Eric Lu, Laurel Johnstone, Michelle Wei, Shudong Sun, Brian Hallmark, Joseph C. Watkins, Hao Helen Zhang, Guang Yao, Floyd H. Chilton

**Affiliations:** School of Nutritional Sciences and Wellness, College of Agriculture and Life Sciences, University of Arizona, Tucson, AZ, USA; Department of Molecular and Cellular Biology, University of Arizona, Tucson, AZ, USA; Department of Mathematics, University of Arizona, Tucson, AZ, USA; Statistics Interdisciplinary Program, University of Arizona, Tucson, AZ, USA; BIO5 Institute, University of Arizona, Tucson, AZ, USA; Center for Precision Nutrition and Wellness, University of Arizona, Tucson, AZ, USA

**Keywords:** sPLA_2_-IIA, Genetic polymorphism, SNP, eQTL, Transcription factor, ChIP-seq-peaks, Transcription factor binding affinity

## Abstract

The secreted phospholipase A_2_ (sPLA_2_) isoform, sPLA_2_-IIA, has been implicated in a variety of diseases and conditions, including bacteremia, cardiovascular disease, COVID-19, sepsis, adult respiratory distress syndrome, and certain cancers. Given its significant role in these conditions, understanding the regulatory mechanisms impacting its levels is crucial. Genome-wide association studies (GWAS) have identified several single nucleotide polymorphisms (SNPs), including rs11573156, that are associated with circulating levels of sPLA_2_-IIA. Through Genotype-Tissue Expression (GTEx), 234 expression quantitative trait loci (eQTLs) were identified for the gene that encodes for sPLA_2_-IIA, *PLA2G2A*. SNP2TFBS (https://ccg.epfl.ch/snp2tfbs/) was utilized to ascertain the binding affinities between transcription factors (TFs) to both the reference and alternative alleles of identified SNPs. Subsequently, ChIP-seq peaks highlighted the TF combinations that specifically bind to the SNP, rs11573156. SP1 emerged as a significant TF/SNP pair in liver cells, with rs11573156/SP1 interaction being most prominent in liver, prostate, ovary, and adipose tissues. Further analysis revealed that the upregulation of PLA2G2A transcript levels through the rs11573156 variant was affected by tissue SP1 protein levels. By leveraging an ordinary differential equation, structured upon Michaelis-Menten enzyme kinetics assumptions, we modeled the PLA2G2A transcription’s dependence on SP1 protein levels, incorporating the SNP’s influence. Collectively, these data strongly suggest that the binding affinity differences of SP1 for the different rs11573156 alleles can influence *PLA2G2A* expression. This, in turn, can modulate sPLA2-IIA levels, impacting a wide range of human diseases.

## Introduction

Members of the secreted phospholipase A_2_ (sPLA_2_) family of enzymes play important roles in numerous biological processes, chiefly by cleaving the *sn*-2 position of glycerophospholipids. This hydrolysis results in an unsaturated fatty acid (UFA) and a 2-lyso-phospholipid. In humans, the 11 isoforms (PLA_2_-IB, PLA_2_-IIA, PLA_2_-IIC, PLA_2_-IID, PLA_2_-IIE, PLA_2_-IIF, PLA_2_-III, PLA_2_-V, PLA_2_-X, PLA_2_-XIIA, and PLA_2_-XIIB) have distinct cellular distributions, enzymatic specificities, and preferences for different polar head groups and UFAs at the sn-3 and sn-2 positions, respectively [1-4]. Collectively, sPLA_2_ isoforms participate in multiple physiological and pathophysiological activities, generating pro- and anti-inflammatory signaling molecules. Their varied functions also encompass remodeling of membrane phospholipids, food phospholipid degradation, and the hydrolysis of bacterial membranes. Notably, the sPLA_2_-IIA isoform, encoded by the *PLA2G2A* gene, efficiently hydrolyzes bacterial membrane phospholipids. This is especially the case for gram-positive bacteria even at minimal sPLA_2_-IIA concentrations, having an essential anti-microbial role in innate immunity [5-8].

Further, circulating sPLA_2_-IIA levels can be reliably measured and have been identified as key biomarkers indicative of the severity or susceptibility linked with cardiovascular disease [9], rheumatoid arthritis [10, 11], cancer [12-15], sepsis [16], and viral infections including Human Hepatitis Virus B [17] and SARS-CoV-2 [18]. Notably, experimental and clinical evidence show that sPLA_2_-IIA plays a critical role in hypotension and multiple organ failure observed in sepsis, COVID-19, and respiratory distress syndrome [19-21]. While numerous genetic variants influencing mRNA levels or blood sPLA_2_-IIA protein levels have been identified, the precise molecular mechanism by which these variants impact sPLA_2_-IIA levels remains to be elucidated. Therefore, gaining insight into the genetic and molecular mechanisms that regulate circulating and tissue sPLA_2_-IIA levels is crucial, as this knowledge can identify populations and individuals vulnerable to either reduced or excessive production of this pivotal enzyme.

The single nucleotide polymorphism (SNP) rs11573156, where G is defined as the reference allele, and C is defined to be an alternative allele in the coding strand sequence, has been strongly associated with circulating levels of sPLA2-IIA. This association is supported by five studies that reveal a dose-dependent allele association between circulating levels and activity [22]. For instance, sPLA2-IIA levels increased by 57% to 62% in those with the homozygous alternative allele (CC) compared to the homozygous reference (GG).

Recognizing the significance of SNPs like rs11573156, our study aimed to understand the genetic and molecular foundations of *PLA2G2A* transcription. We focused on examining transcription factor (TF) binding sites where PLA2G2A-eQTLs are located and how different alleles at rs11573156 influence TF binding affinity. We then constructed an ordinary equation model, assuming Michaelis-Menten kinetics, to predict the molecular mechanism of SP1 activating *PLA2G2A* transcription. Within this model, the difference between the alternative allele and the reference allele is expressed in the value of the Michaelis constant, *K*. This model was tested with data of SP1 protein and *PLA2G2A* transcripts for each tissue obtained from GTEx and Human Protein Atlas (HPA), respectively. Our findings reveal that rs11573156 affects *PLA2G2A* transcription via the differential binding affinities of the two alleles to the SP1 TF, and this likely constitutes a primary genetic/molecular mechanism affecting circulating sPLA2-IIA levels.

## Materials and Methods

### Collection of PLA2G2A eQTLs in different tissues

The GTEx Portal V8 (https://gtexportal.org) [43] was utilized to determine *PLA2G2A* expression quantitative trait loci (eQTLs) across 18 tissues including thyroid, testis, stomach, skin-sun exposed (lower leg), skin-not sun exposed (suprapubic), prostate, ovary, nerve-tibial, muscle-skeletal, lung, liver, heart-left ventricle, heart-atrial appendage, esophagus-muscularis, esophagus-gastroesophageal junction, artery-aorta, adipose-visceral (omentum), and adipose-subcutaneous. GTEx includes data measured in tissues collected within 24 hours of death for 948 individuals (32.9% female; 84.6% white, 12.9% African American, 1.3% Asian, 1.1% unknown).

### TF binding to alleles of predicted eQTLs

The SNP2TFBS web interface (http://ccg.vital-it.ch/snp2tfbs/) was used to determine which eQTL SNPs were predicted to affect single or multiple TFs. SNP2TFBS is a comprehensive database of regulatory SNPs affecting predicted transcription factor binding site (TFBS) affinity [29]. A SNP’s effect on TF binding is estimated based on a position weight matrix (PWM) model for the binding specificity of the corresponding factor, which are fixed length TFBS models represented by a matrix of probabilities reflecting the occurrence frequencies of bases at binding site positions.

SNP2TFBS uses the JASPAR database (http://jaspar.genereg.net) as a source of the TF binding profiles. JASPAR provides PWMs as base frequency matrices. SNP2TFBS reports raw binding scores, which are computed as the sum of the position-specific weights over all bases of the binding site. It uses SNP data from the 1000 Genomes Project (minor allele frequency > 0.001), human genome assembly GRCh37/hg19, and RefSeq gene annotations provided by ANNOVAR [44]. In this study, eQTLs corresponding to upregulated or downregulated *PLA2G2A* gene expression were queried with SNP2TFBS to obtain a list of predicted TFs corresponding to those SNPs.

### Verification of the predicted transcription factors by ChIP seq- peaks analysis

ChIP-seq Peaks (340 factors in 129 cell types) from ENCODE 3, as downloaded from the UCSC Genome Browser alignment to GRCh38/hg38, were utilized to examine whether the candidate TFs have binding sites. This data set represents a comprehensive set of human transcription factor binding sites based on ChIP-seq experiments generated by production groups in the ENCODE Consortium between February 2011 and November 2018. If candidate TFs from SNP2TFBS had binding sites within the PLA2G2A eQTLs region (chromosome1; 18990581-20971970), we checked whether the corresponding SNP was within that TF binding site.

### Gene expression data normalized by transcript per million (TPM)

The RNA-sequencing (RNA-seq) data (Gene TPMs) were obtained from the GTEx portal open access database (https://gtexportal.org) on 04/21/2022. The genotyping data was obtained after approval of a dbGaP application and downloaded as dbGaP accession number phg001219.v1 on 10/02/2022.

### SP1 protein expression in different tissues

SP1 protein expression data by tissue are publicly available from the Human Protein Atlas’s website (https://www.proteinatlas.org/about/download). Expression values are semiquantitative, derived from immunohistochemistry of tissue micro arrays, and take on 4 possible values: “Not Detected”, “Low”, “Medium”, and “High”. Tissues not present in GTEx were omitted. When a tissue had multiple recorded cell types, the protein expression profile with the largest proportion of cell type within a tissue was selected.

### Modeling the correlation between SP1 levels and *PLA2G2A* gene expression

Two models were utilized for correlation analysis of SP1 levels and *PLA2G2A* expression. Initially, a simple linear regression analysis was employed applying TFs as the independent variable and *PLA2G2A* expression as the dependent variable. The coefficient of determination (R^2^) was calculated to compare across several TFs.

A second model utilized the simple ordinary differential equation shown below to provide a mathematical framework for these observations.

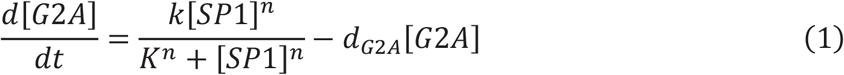

The structure of the equation was based on standard Michaelis-Menten enzyme kinetics assumptions.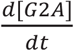 describes the rate of the change of *PLA2G2A* transcript concentration over time. *k* is a constant representing protein-dependent PLA2G2A transcription. *K* is the Michaelis constant. *n* is the hill coefficient for cooperativity. *d*_*G*2*A*_is the degradation constant of *PLA2G2A* transcripts. The steady state concentration of *PLA2G2A* expression was solved (1) to yield the following equation:

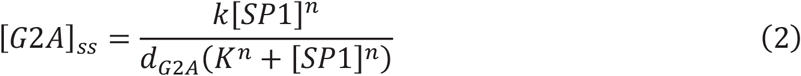

Parameters were loosely selected to fit the observed trend and are listed as follows: *k* =1, *d*_[*G*2*A*]_ =0.01, *n*=2, *K*_*C*_=1, and *K*_*G*_=5 where *K*_*C*_ and *K*_*G*_ represent the Michaelis constants for the CC, and GG + GC genotypes, respectively. *PLA2G2A* steady state transcript concentrations then were simulated over a range of SP1 protein concentrations from 0 – 20 arbitrary units.

## Results

### eQTLs of *PLA2G2A* in Human Tissues

To investigate the potential genetic basis for variations in *PLA2G2A* expression in humans, the relationship between the SNPs in the human genome and *PLA2G2A* expression levels were examined. Specifically, we leveraged data from the Genotype-Tissue Expression (GTEx) project, which correlates genome-wide SNPs with variations in gene expression across multiple human tissues. Our analysis identified 234 eQTLs significantly associated with *PLA2G2A* expression (with a false discovery rate FDR ≤0.05) in GTEx (V8) (**Figure 1A**). Located on chromosome 1 (hg38; chr1: 19975431-19980416), these eQTLs primarily occupy non-coding regions proximal to and flanking the *PLA2G2A* gene locus. Of the 234 eQTLs, 171 are associated with increased *PLA2G2A* mRNA expression, while 63 are linked with its downregulation. Interestingly, 37 of these eQTLs were observed in two or more tissues, with each displaying a consistent effect on *PLA2G2A* expression across these tissues, either upregulating or downregulating its levels (**Figure 1B, Supplementary Table 1**). This consistency pointed to the potential roles these 37 eQTLs play in modulating *PLA2G2A* expression throughout the human body. Consequently, our subsequent analysis was centered on these 37 eQTLs.

**Figure 1.**
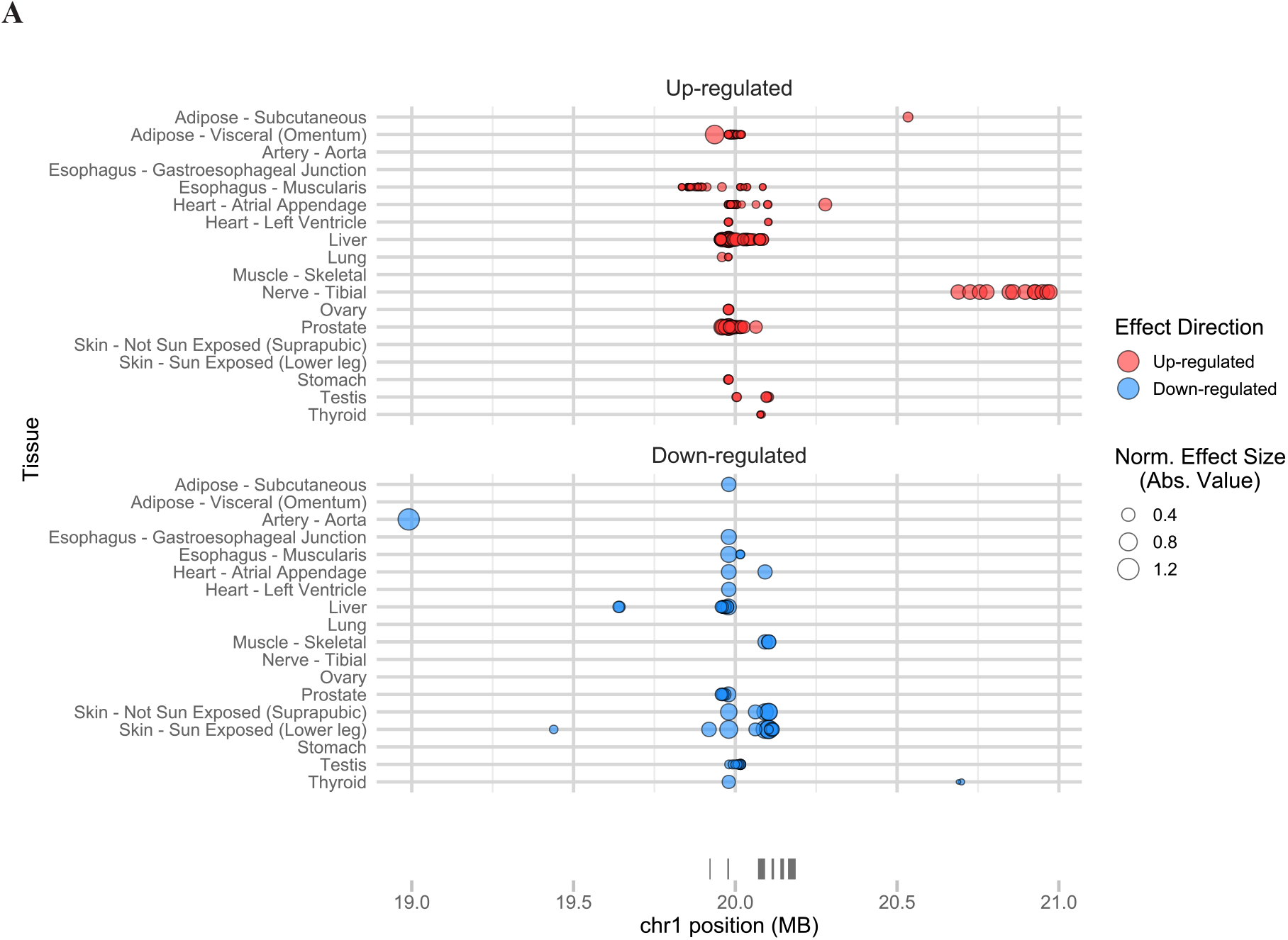
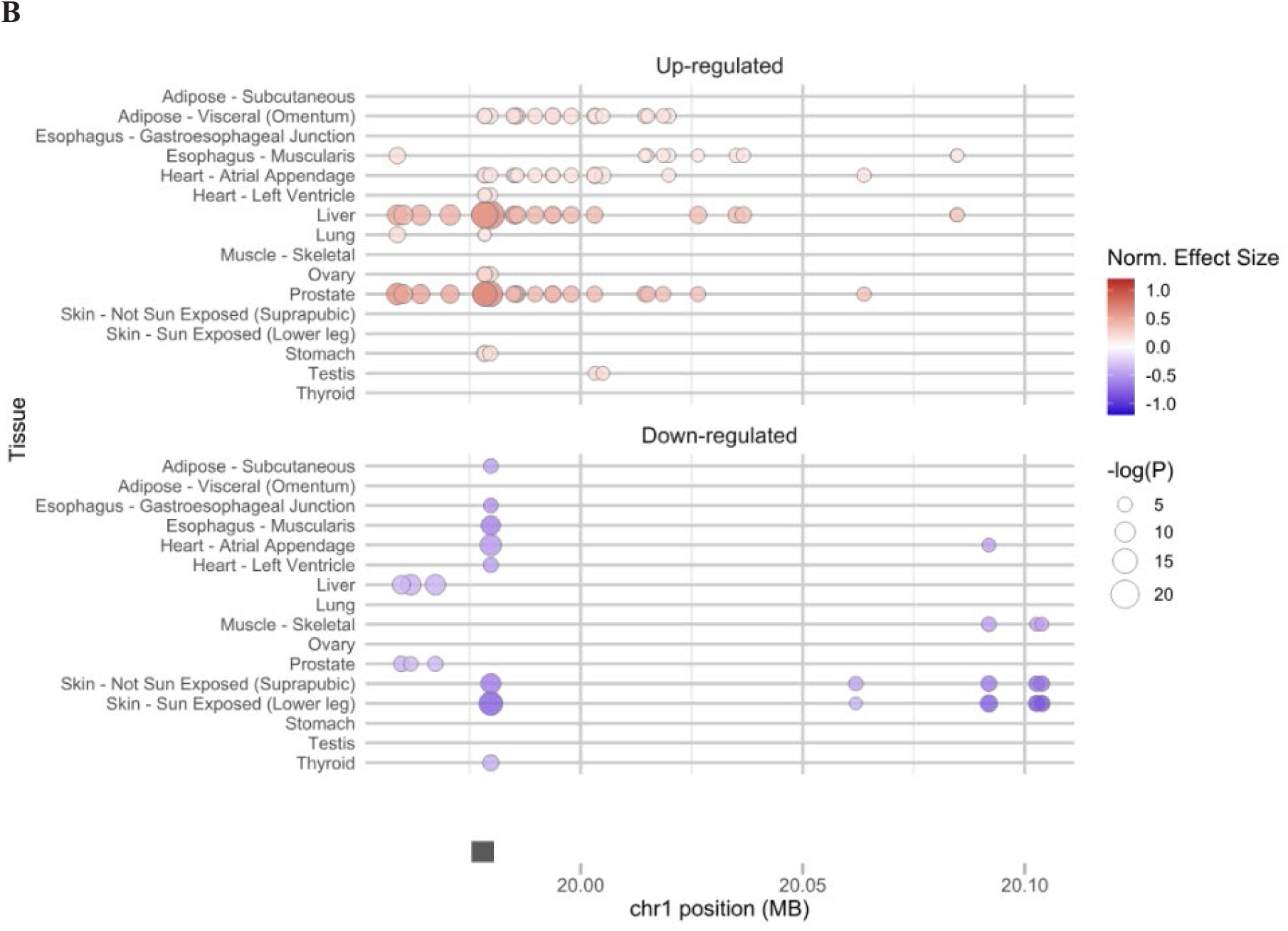
A) The location of 234 eQTLs for *PLA2G2A*. X-axis shows the chromosomal position, and Y axis shows tissues. The marker size indicates the magnitude of the effect size. The marker color indicates the effect direction in alternative alleles compared to reference alleles. B) The location of 37 eQTLs shared among more than one tissue. X-axis shows the chromosomal position, and Y axis shows tissues. The marker size indicates the magnitude of the negative logarithm of the P-value. The color intensity indicates the magnitude of normalized effect size. The color indicate the effect direction in alternative alleles compared to reference alleles. Red: up-regulated, Blue: down-regulated. The normalized effect size (NES) of the eQTLs is defined as the slope of the linear regression and is computed as the effect of the alternative allele (ALT) relative to the reference allele (REF) in the human genome reference GRCh38/hg38, as described in GTEx Portal (https://gtexportal.org/home/methods).

### Identification of eQTL-Bound Transcription Factors

eQTLs have the potential to modulate gene expression by various mechanisms, including transcription, RNA processing, RNA stability, and translation [23-25]. One of the predominant mechanisms involves the alteration of transcription factor (TF) binding sites on the targeted gene promoters [26-28]. To identify the TF binding sites that may be affected by *PLA2G2A* eQTLs, we examined the 37 eQTLs (**Figure 1B, Supplementary Table 1**) using SNP2TFBS, a computer program that predicts how a SNP may affect TF binding site sequence and binding affinity [29].

From the SNP2TFBS analysis, 9 of the 37 eQTLs were associated with TF binding sites, Notably, 7 of these 9 eQTLs had the potential to influence TF binding affinities due to sequence variations between the reference and alternative SNP alleles (**Table 1**). In particular, 7 eQTLs were predicted to affect the binding of 8 TFs: EBF1, ZNF263, SP1, PPARG_RXRA, SREBF1, SREBF2, Gata1, and Foxq1 (**Table 1**).

**Table 1:** Candidate functional variants associated with PLA2G2A gene expression determined by SNP2TFBS. ***Ref*** reference allele, ***Alt*** alternative allele, ***Effect*** the direction of the eQTL’s effect for the PLA2G2A gene expression of alternative allele compared to reference allele, ***eQTL tissues*** tissue distribution of each eQTL, ***TF*** transcription factor.

| SNP (eQTL) rsID | Ref | Alt | Effect | eQTL tissues | Matched TFs |
| --- | --- | --- | --- | --- | --- |
| rs12733637 | A | G | down | Liver, Prostate | EBF1 |
| rs1891320 | C | T | up | Liver, Prostate | ZNF263 |
| rs11573156 | G | C | up | Liver, Prostate, Ovary, Adipose-Visceral, Stomach, Heart-Atrial Appendage, Heart-Left Ventricle | SP1 |
| rs1796923 | T | G | down | Skin - Sun Exposed (Lower leg), Heart-Atrial Appendage, Skin-Not Sun Exposed (Suprapubic), Esophagus-Muscularis, Thyroid, Heart-Left Ventricle, Adipose-Subcutaneous, Esophagus-Gastroesophageal Junction | PPARG_RXRA |
| rs114156017 | C | T | up | Prostate, Adipose-Visceral, Heart-Atrial Appendage | SREBF1, SREBF2 |
| rs149238894 | G | A | up | Prostate, Adipose-Visceral, Heart-Atrial Appendage | Gata1 |
| rs139466428 | TG | T | up | Prostate, Adipose-Visceral, Esophagus-Muscularis | Foxq1 |

### Examination of TF-eQTL Physical Binding

The relationship between eQTLs and TFs in the tissues they influence were next examined. We posited that if an eQTL modulates gene expression (in this case PLA2G2A) by affecting TF binding affinity, there would be TF binding to the eQTL site in the tissues where the eQTL has an effect. To address this question, we utilized the ENCODE chromatin immunoprecipitation (ChIP)-seq dataset, which maps the chromatin binding patterns of 340 TFs across 129 cell lines or tissue types. From the the 8 TFs hypothesized to be influenced by *PLA2G2A* eQTLs, 6 TFs were evaluated in ENCODE dataset (**Table 2**). However, only one TF, SP1, had been studied in a primary tissue, specifically, the liver.

**Table 2:** The list of TFs in ENCODE (ChIP-seq peaks database) in the current analysis.

| TF | Cell line | Cell line origin |
| --- | --- | --- |
| EBF1 | GM12878 | B-lymphocyte |
| ZNF263 | HEK293 | Kidney |
| SP1 | A549 | Lung |
|  | H1-hESC | Embryo |
|  | HEK293T | Kidney |
|  | HepG2 | Liver |
|  | K562 | Bone marrow |
|  | liver | Liver |
|  | MCF-7 | Breast |
| GATA1 | erythroblast | erythroblast |
|  | K562 | Bone marrow |
| SREBF1 | A549 | Lung |
|  | K562 | Bone marrow |
|  | MCF-7 | Breast |
| SREBF2 | A549 | Lung |
|  | HeLa-S3 | Uterus cervix |

The ENCODE data revealed that SP1 binds to the chromatin region of the eQTL rs11573156 **(Figure 2**). While the other 7 TFs predicted by SNP2TFBS may also interact with the *PLA2G2A* eQTLs, the relationship between SP1 and rs11573156 in modulating *PLA2G2A* expression became a primary focus. This decision was based on two main factors: 1) The binding of SP1 to this eQTL was validated by the ENCODE ChIP-seq data from liver samples, and 2) Among all tissues explored in GTEx, the liver displayed the most pronounced eQTL effect of rs11573156 on *PLA2G2A* expression (**Supplementary Table 2**).

**Figure 2:**
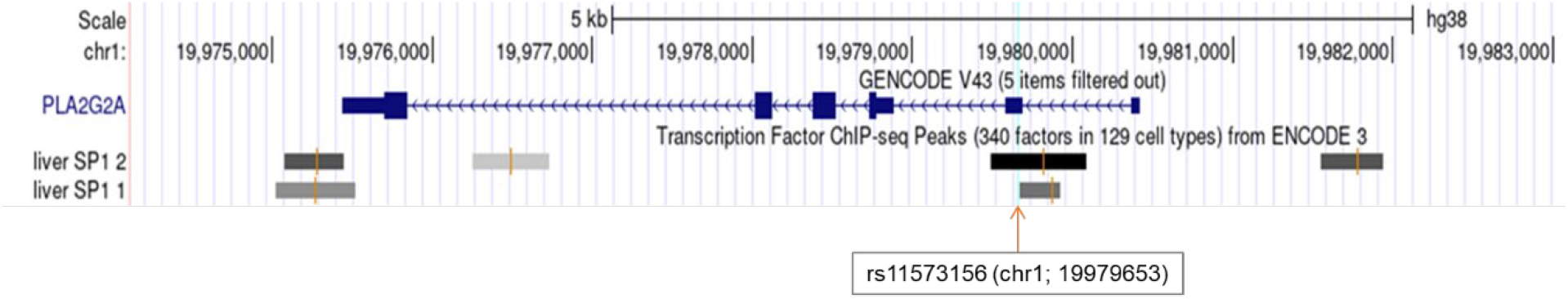
Display of the TF binding sites and the location of SNPs by UCSC genome browser. The figure zoomed in on the range where the SNP of rs11573156 is present. Transcription Factor (TF) ChIP-seq Peaks (340 factors in 129 cell types) from ENCODE 3 was used to identify TF binding sites (sequences). This database provides TF binding sites determined by chromatin immunoprecipitation combining DNA sequencing using each cell line. The gray and black rectangles indicate the respective TF-binding sequences in each liver cell derived from separate individual. A vertical light blue line indicates the location of SNP of rs11573156.

### SP1 Bound to rs11573156 as a Transcriptional Activator of *PLA2G2A*

If SP1 binds to the eQTL rs11573156 in the liver and thereby influences *PLA2G2A* expression, a discernible correlation between liver SP1 and *PLA2G2A* would be anticipated. Specifically, considering the higher binding affinity of SP1 to the alternative allele (C) of rs11573156 over the reference allele (G) (as shown in **Table 1**) coupled with the positive effect of the alternative allele C on *PLA2G2A* expression, we postulated that there would be a positive correlation between SP1 and *PLA2G2A* levels in the liver. Indeed, in the GTEx dataset containing 208 liver donor samples, SP1 was positively associated with *PLA2G2A* expression (correlation coefficient, r = 0.31; **Figure 3A**). This was consistent with the role of SP1 as a transcriptional activator of *PLA2G2A* gene in the liver.

**Figure 3.**
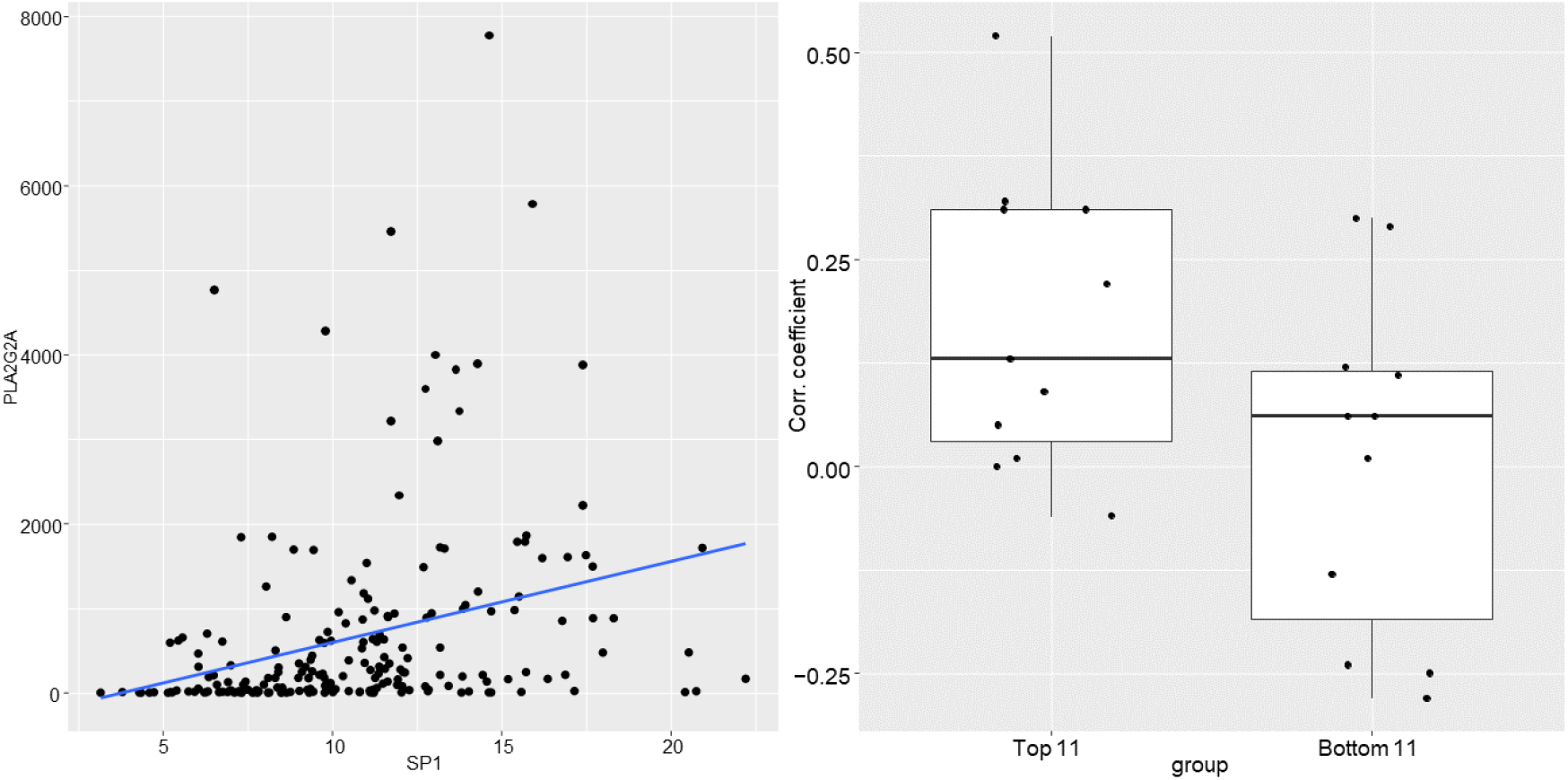
A) Correlation analysis of the gene expression between PLA2G2A and SP1. X-axis shows transcripts per million (TPM) of SP1 and Y-axis shows TPM of PLA2G2A. B) Box plots of the correlation efficient between SP1 and PLA2G2A TPMs in each tissue. Top11; top 11 tissues by effect size of the eQTL of rs11573156. Bottom 11; bottom 11 tissues by effect size.

We next focused on the top 11 tissues or tissue subtypes from the GTEx dataset (out of a total of 48) where the effect of eQTL rs11573156 on PLA2G2A expression was most pronounced. These tissues were defined by a single-tissue p-value < 0.001 and the multi-tissue posterior probability m-value = 1 (**Supplementary Table 2**). We hypothesized that if SP1 binds to the eQTL rs11573156 and elevates *PLA2G2A* expression, then the correlation between SP1 with *PLA2G2A* would be stronger in these 11 tissues compared to others This expectation was validated in our analysis. As illustrated in **Figure 3B**, the correlation coefficients between SP1 and PLA2G2A in these top 11 tissues (with an m-value of 1) were significantly higher (p = 0.03) than in the control tissues (bottom 11 with m-value < 0.21; refer to **Supplementary Table 2**). Although a positive correlation between SP1 and *PLA2G2A* levels isn’t definitive proof of a functional causal relationship, these data are consistent with the interpretation that in the majority (if not all) of the top 11 tissues, SP1 binds to the eQTL rs11573156, thereby affecting (transactivating) *PLA2G2A* expression.

### SP1 Differential Binding to rs11573156 Alleles Influences *PLA2G2A* Expression Variation

In the analysis of the top 11 tissues, there was a striking variation in the effect size of the eQTL rs11573156, with fluctuations spanning over a 5-fold range (from 0.12 to 0.68). Given that the alternative allele of rs11573156 is associated with higher SP1 binding affinity, these findings raised the question of why does this mechanism exhibit such pronounced variability in effect sizes across distinct tissues? To explore the molecular complexities of how SP1 transcriptionally activates *PLA2G2A*, we developed a basic ordinary differential equation model based on the premise of Michaelis-Menten kinetics:

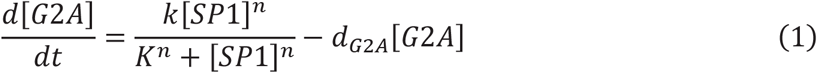

In this model, the change of *PLA2G2A* transcript level over time 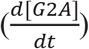 is a function of SP1 (transcription factor) protein level (with *k, K, n* and *d*_*G*2*A*_ being constants, see Method for details). The difference between the alternative allele (C) and the reference allele (G) of rs11573156 is reflected in the value of the Michaelis constant *K* that is inversely correlated with SP1 binding affinity (*K*_*C*_ = 1, representing a higher SP1 binding affinity to the alternative allele C; *K*_*G*_ = 5, representing a lower SP1 binding affinity to the reference allele G).

This model suggests that because of the difference in *K*_*C*_ vs. *K*_*G*_, the steady-state level of *PLA2G2A* transcripts [G2A]_ss_ is higher with the alternative allele (C) than with the reference allele (G). Importantly, the model predicts that the discrepancy in [G2A]_ss_ between the two alleles, represented as Δ[G2A]_ss_, becomes more pronounced at lower concentrations of SP1 protein levels. This divergence then gradually diminishes as SP1 concentrations rise (as depicted in **Figure 4A**). Thus, this result can largely be explained by the “saturating” effect at high SP1 levels, where the excess amount of SP1 protein has the capacity to fully transactivate PLA2G2A even to the lower binding affinity allele of rs11573156.

**Figure 4:**
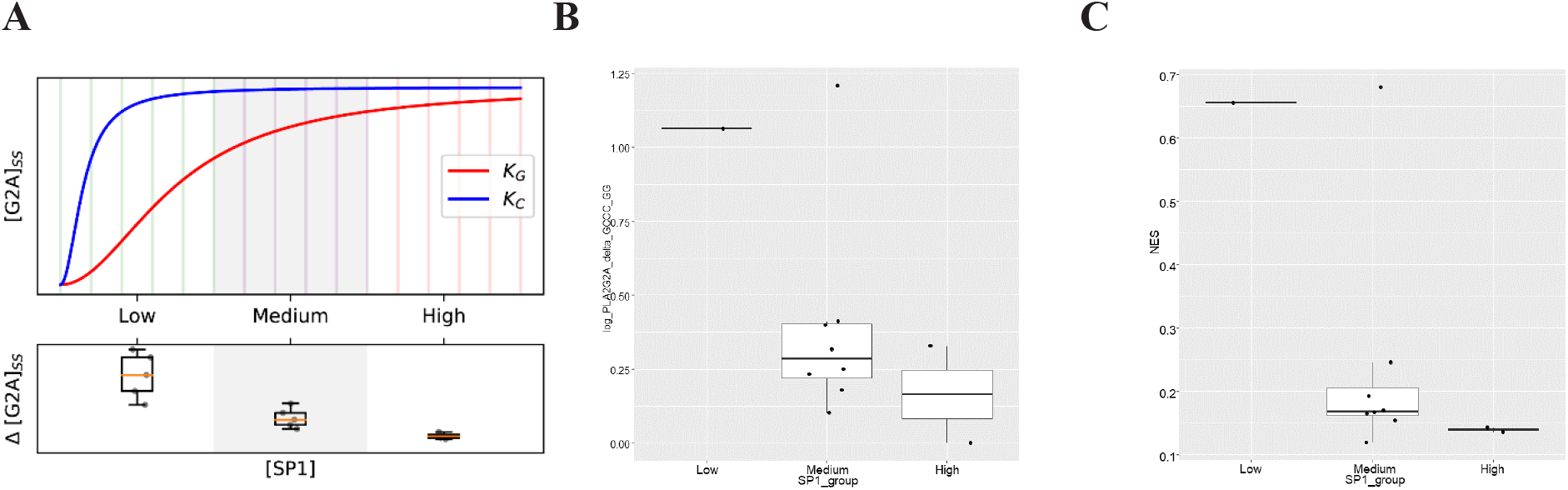
**A**) Modeled *PLAG2A* (G2A) allele expression. In the upper panel, the ODE across an arbitrary range of [SP1] was simulated using an arbitrary parameter set to get G2A expression levels. The lower panel boxplots contained 15 points total, 5 per box, and 1 per bin. The Δ[*G*2*A*]_*ss*_ was the subtraction of the C and G allele PLA2G2A gene expression levels and 5 bins were created for each Low, Medium, and High [SP1]. The exact parameters are *range* ([*SP*1]) = [0, 20], *k* = 1;*d*_*G*2*A*_ = 0.01;*nn* = 2;*K*_*C*_ = 1;*K*_*G*_ = 5. **B**) Empirical delta PLA2G2A allele expression. The differences (delta logTPM) of PLA2G2A transcript levels by genotype were calculated in every tissue, and categorized into three groups depending on SP1 protein levels, low, medium, and high, and box plotted them. t-test was performed within tissue between the two alleles to display statistical significance in G2A expression between the two alleles. GG was designated as G, and GC and CC as C. **C**) Normal effect size in each tissue categorized into the same three groups as Figure 4**B**.

### Validation of the Model Predictions Using SP1 Protein Levels Across Tissues

To test the model predictions, we obtained the SP1 protein levels across diverse tissues from the Human Protein Atlas (HPA). SP1 protein levels were emphasized for a couple of reasons: 1) It’s the protein level (rather than the mRNA level) of the transcription factor SP1 that impacts *PLA2G2A* expression, and 2) the SP1 mRNA levels across different tissues do not always correlate well with SP1 protein levels, due to potential tissue-specific differences in posttranscriptional modifications, translation, and protein stability, etc.

HPA estimates protein levels in different human tissues in three categories, low, medium, and high. The categorical SP1 protein levels in the top 11 tissues are shown in Suppl. Table 2. When the *PLA2G2A* transcript level differential (ΔlogTPM) between the C and G alleles in each of the 11 tissues was plotted against their corresponding SP1 protein categories, a clear trend emerged. This trend was consistent with the model’s predictions: the difference in *PLA2G2A* transcript levels between alleles C and G was most significant at the lowest SP1 protein level, tapering off as SP1 levels ramped up to medium and then high (**Figure 4B**). Similarly, the effect size of the eQTL rs11573156, which is positively correlated with the *PLA2G2A* level difference between the C and G alleles, also exhibits a similar trend that is inversely correlated with SP1 protein level in the tissue (Fig. 4C). Cumulative, these results strongly support the hypothesis that variations in *PLA2G2A* gene expression associated with the eQTL rs11573156 alleles are significantly, and inversely, influenced by the tissue-specific SP1 protein levels.

## Discussion

Much of the early work with the sPLA_2_-IIA isoform centered on its bactericidal role where it was shown to increase 100 to 1000-fold (up to 5 μg/ml) in serum/plasma during infectious sepsis [30]. However, studies beginning in the mid-1980s, also identified its capacity at persistently high concentrations to induce lethal multiple organ failure associated with sepsis [18, 20, 31]. In fact, Nyman and colleagues demonstrated that sPLA_2_-IIA levels could effectively predicted lethal multiple organ failure when measured at later time points (>3 days in the ICU) in sepsis patients [20].

More recently, our group discovered that deceased COVID-19 patients, primarily those succumbing to lethal multiple organ failure, also exhibited considerably elevated levels of circulating, catalytically active sPLA_2_-IIA [18]. Elevated sPLA_2_-IIA levels paralleled several indices of COVID-19 disease severity such as kidney dysfunction, hypoxia, multiple organ dysfunction. Moreover, three machine learning models identified sPLA_2_-IIA levels as the pivotal factor in differentiating patients who died from sepsis due to COVID-19. A pressing question emerging from these studies and other clinical conditions, such as cardiovascular disease [9], rheumatoid arthritis [10], and cancer [12-15], is: Why do only certain patients produce and release high concentrations of sPLA2-IIA from tissues, especially the liver?

Several GWASs have demonstrated the impact of genetic variation on sPLA_2_-IIA levels and further revealed that rs11573156 G>C polymorphism is highly associated with circulating level of sPLA_2_-IIA [32-35]. The allele frequencies mean for all global populations are 77% for the G reference allele and 23% for the C alternative allele. Frequencies of the C allele varying from 5% (Asian) to 36% (Estonian) in different populations.

The current study demonstrated that among the 234 eQTLs for *PLA2G2A*, rs11573156 in the liver showed the most dramatic changes in the gene expression levels. This SNP exclusively affects *PLA2G2A* gene expression, and it is not identified as an eQTL with any of the other 11 sPLA2 genes, including the 6 (*PLA2G2E, PLA2GA, PLA2G5, PLA2G2D, PLA2GF, and PLA2G2C*) that occur as a gene cluster (hg38 chr1:19,920,009-20,186,518) on chromosome 1 (data not shown). Furthermore, the pQTL (protein quantitative trait locus) analysis also indicated that rs11573156 is strongly associated with protein levels of sPLA_2_-IIA [33-36].

The rs11573156 SNP is located in the 5’ untranslated region (5’-UTR) of the *PLA2G2A* gene, acting as a TF binding site that regulates mRNA expression [37]. The current study revealed this region containing rs11573156 is a crucial SP1 binding site that modulates *PLA2G2A* transcript levels. Specifically, in tissues such as the liver where levels of SP1 are limited (not saturated), there is a pronounced difference in the binding affinity of SP1 to the G allele versus the C allele. Moreover, there appears to be a direct relationship between *PLA2G2A* transcript levels and sPLA_2_-IIA protein expression, as highlighted by the fact that rs11573156 serves as a pQTL. Notably, the primary organ thought to produce sPLA_2_-IIA released into circulation is the liver [38-40]. This aligns with the our observation that rs11573156 alleles via their binding affinity to SP1 modulated differences in *PLA2G2A* expression in the liver and sPLA2-IIA protein levels in circulation.

This study has important limitations. First, ChIP seq-peaks analysis does not include all potential tissues where sPLA_2_-IIA may play a biological role. The alleles at rs11573156 are associated with significant *PLA2G2A* gene expression differences in the liver, prostate, ovary, stomach, adipose-visceral, heart-left ventricle, and heart-atrial appendage. However, given ChIP-seq data were only available in a liver cell line, it remains unclear whether SP1-mediated *PLA2G2A* regulation occurs in these other seven tissues. Second, the methodology employed in this study solely investigated the affinity of SNP TF binding sites as a means to regulate *PLA2G2A* expression. It’s worth noting that some SNPs might also impact a variety of other mechanisms, including mRNA stability, subcellular localization, and the modulation of micro RNAs (miRNAs) [41, 42].

Given the diverse functions and diseases where sPLA*2*-IIA plays a modulatory role, it is imperative to determine how levels of this pivotal enzyme are controlled and whether its synthesis can be inhibited. This paper describes an allele-specific transcription factor binding interaction between rs11573156 (the SNP that has been most associated sPLA2-IIA levels) and SP1. This interaction appears to regulate *PLA2G2A* expression in the liver. Unraveling a genetic molecular mechanism that controls sPLA2-IIA levels in the primary organ where it is generated is a significant advancement in this field. Furthermore, studies like this pave the way for precision medicine approaches that can modulate *PLA2G2A* expression by altering the dynamics of SP1 binding to SNPs, such as rs11573156, especially in potentially fatal diseases like infectious sepsis.

## Conflict of Interest

Dr. Chilton is a cofounder of Tyrian Omega, Inc, and Resonance Pharma, Inc. These companies use microorganisms to generate omaga-3 fatty acids and develop diagnostic and therapeutics for lipid targets, respectively. These commercial relationships are managed by the Office for Responsible Outside Interests at University of Arizona.

## Author Contributions

FHC and GY conceived and designed the study. AH, EL, SS, LJ, MW, BH, HZ, and GY did the statistical analysis, modeling, and visualization. AH and FHC wrote the initial manuscript draft. The discussion was developed and written by FHC, AH, BH, JCW, HZ, and GY. All authors contributed to revisions.

## Acknowledgment

This work was supported by National Center for Complementary and Integrative Health [R01 AT008621; FHC] and U.S. Department of Agriculture [ARZT-1361680-H23-157; FHC].

---

**Supplementary Table 1**: The list of eQTLs for *PLA2G2A*. Significant single tissue eQTLs for PLA2G2A (ENSG00000188257.10) were collected from the GTEx database and listed by SNP rsID with tissues identified. eQTLs do not have the same effect on all tissues and are counted based on the combination of tissue and SNP. 234 eQTLs were detected as significant eQTLs for *PLA2G2A* in total. The information of functional consequence and variant type (2^nd^ and 3^rd^ column) were obtained from the Single Nucleotide Polymorphism Database (dbSNP) hosted by National Human Center for Biotechnology Information (NCBI) (https://www.ncbi.nlm.nih.gov/snp/).

**Supplementary Table 2:** Multi-tissue comparison of the Normalized effect size (NES) of rs11573156 C allele (Alternative) against G allele (Reference), combined with Human Protein Atlas (HPA) protein levels in top 11 NES tissues.

## References

[1] Burke JE, Dennis EA. Phospholipase A2 structure/function, mechanism, and signaling. J Lipid Res. 2009;50 Suppl:S237–42.

[2] Cupillard L, Koumanov K, Mattei MG, Lazdunski M, Lambeau G. Cloning, chromosomal mapping, and expression of a novel human secretory phospholipase A2. J Biol Chem. 1997;272:15745–52.

[3] Murakami M, Sato H, Miki Y, Yamamoto K, Taketomi Y. A new era of secreted phospholipase A(2). J Lipid Res. 2015;56:1248–61.

[4] Six DA, Dennis EA. The expanding superfamily of phospholipase A(2) enzymes: classification and characterization. Biochim Biophys Acta. 2000;1488:1–19.

[5] Dore E, Boilard E. Roles of secreted phospholipase A(2) group IIA in inflammation and host defense. Biochim Biophys Acta Mol Cell Biol Lipids. 2019;1864:789–802.

[6] Gronroos JO, Laine VJ, Nevalainen TJ. Bactericidal group IIA phospholipase A2 in serum of patients with bacterial infections. J Infect Dis. 2002;185:1767–72.

[7] Movert E, Wu Y, Lambeau G, Kahn F, Touqui L, Areschoug T. Secreted group IIA phospholipase A2 protects humans against the group B streptococcus: experimental and clinical evidence. J Infect Dis. 2013;208:2025–35.

[8] van Hensbergen VP, Wu Y, van Sorge NM, Touqui L. Type IIA Secreted Phospholipase A2 in Host Defense against Bacterial Infections. Trends Immunol. 2020;41:313–26.

[9] Kugiyama K, Ota Y, Takazoe K, Moriyama Y, Kawano H, Miyao Y, et al. Circulating levels of secretory type II phospholipase A(2) predict coronary events in patients with coronary artery disease. Circulation. 1999;100:1280–4.

[10] Liu NJ, Chapman R, Lin Y, Mmesi J, Bentham A, Tyreman M, et al. Point of care testing of phospholipase A2 group IIA for serological diagnosis of rheumatoid arthritis. Nanoscale. 2016;8:4482–5.

[11] Seilhamer JJ, Pruzanski W, Vadas P, Plant S, Miller JA, Kloss J, et al. Cloning and recombinant expression of phospholipase A2 present in rheumatoid arthritic synovial fluid. J Biol Chem. 1989;264:5335–8.

[12] Dong Z, Liu Y, Scott KF, Levin L, Gaitonde K, Bracken RB, et al. Secretory phospholipase A2-IIa is involved in prostate cancer progression and may potentially serve as a biomarker for prostate cancer. Carcinogenesis. 2010;31:1948–55.

[13] Kupert E, Anderson M, Liu Y, Succop P, Levin L, Wang J, et al. Plasma secretory phospholipase A2-IIa as a potential biomarker for lung cancer in patients with solitary pulmonary nodules. BMC Cancer. 2011;11:513.

[14] Oleksowicz L, Liu Y, Bracken RB, Gaitonde K, Burke B, Succop P, et al. Secretory phospholipase A2-IIa is a target gene of the HER/HER2-elicited pathway and a potential plasma biomarker for poor prognosis of prostate cancer. Prostate. 2012;72:1140–9.

[15] Zhang C, Yu H, Xu H, Yang L. Expression of secreted phospholipase A2-Group IIA correlates with prognosis of gastric adenocarcinoma. Oncol Lett. 2015;10:3050–8.

[16] Tan TL, Goh YY. The role of group IIA secretory phospholipase A2 (sPLA2-IIA) as a biomarker for the diagnosis of sepsis and bacterial infection in adults-A systematic review. PLoS One. 2017;12:e0180554.

[17] Zhu C, Song H, Shen B, Wu L, Liu F, Liu X. Promoting effect of hepatitis B virus on the expressoin of phospholipase A2 group IIA. Lipids Health Dis. 2017;16:5.

[18] Snider JM, You JK, Wang X, Snider AJ, Hallmark B, Zec MM, et al. Group IIA secreted phospholipase A2 is associated with the pathobiology leading to COVID-19 mortality. J Clin Invest. 2021;131.

[19] Guidet B, Piot O, Masliah J, Barakett V, Maury E, Bereziat G, et al. Secretory non-pancreatic phopholipase A2 in severe sepsis: relation to endotoxin, cytokines and thromboxane B2. Infection. 1996;24:103–8.

[20] Nyman KM, Uhl W, Forsstrom J, Buchler M, Beger HG, Nevalainen TJ. Serum phospholipase A2 in patients with multiple organ failure. J Surg Res. 1996;60:7–14.

[21] Vadas P, Pruzanski W, Farewell V. A predictive model for the clearance of soluble phospholipase A2 during septic shock. J Lab Clin Med. 1991;118:471–5.

[22] Holmes MV, Simon T, Exeter HJ, Folkersen L, Asselbergs FW, Guardiola M, et al. Secretory phospholipase A(2)-IIA and cardiovascular disease: a mendelian randomization study. J Am Coll Cardiol. 2013;62:1966–76.

[23] Cano-Gamez E, Trynka G. From GWAS to Function: Using Functional Genomics to Identify the Mechanisms Underlying Complex Diseases. Front Genet. 2020;11:424.

[24] Roadmap Epigenomics C, Kundaje A, Meuleman W, Ernst J, Bilenky M, Yen A, et al. Integrative analysis of 111 reference human epigenomes. Nature. 2015;518:317–30.

[25] Soskic B, Cano-Gamez E, Smyth DJ, Rowan WC, Nakic N, Esparza-Gordillo J, et al. Chromatin activity at GWAS loci identifies T cell states driving complex immune diseases. Nat Genet. 2019;51:1486–93.

[26] Farh KK, Marson A, Zhu J, Kleinewietfeld M, Housley WJ, Beik S, et al. Genetic and epigenetic fine mapping of causal autoimmune disease variants. Nature. 2015;518:337–43.

[27] Ishigaki K. Beyond GWAS: from simple associations to functional insights. Semin Immunopathol. 2022;44:3–14.

[28] Musunuru K, Strong A, Frank-Kamenetsky M, Lee NE, Ahfeldt T, Sachs KV, et al. From noncoding variant to phenotype via SORT1 at the 1p13 cholesterol locus. Nature. 2010;466:714–9.

[29] Kumar S, Ambrosini G, Bucher P. SNP2TFBS - a database of regulatory SNPs affecting predicted transcription factor binding site affinity. Nucleic Acids Res. 2017;45:D139–D44.

[30] Nevalainen TJ, Eerola LI, Rintala E, Laine VJ, Lambeau G, Gelb MH. Time-resolved fluoroimmunoassays of the complete set of secreted phospholipases A2 in human serum. Biochim Biophys Acta. 2005;1733:210–23.

[31] Vadas P. Elevated plasma phospholipase A2 levels: correlation with the hemodynamic and pulmonary changes in gram-negative septic shock. J Lab Clin Med. 1984;104:873–81.

[32] Akinkuolie AO, Lawler PR, Chu AY, Caulfield M, Mu J, Ding B, et al. Group IIA Secretory Phospholipase A(2), Vascular Inflammation, and Incident Cardiovascular Disease. Arterioscler Thromb Vasc Biol. 2019;39:1182–90.

[33] Benson MD, Yang Q, Ngo D, Zhu Y, Shen D, Farrell LA, et al. Genetic Architecture of the Cardiovascular Risk Proteome. Circulation. 2018;137:1158–72.

[34] Wootton PT, Drenos F, Cooper JA, Thompson SR, Stephens JW, Hurt-Camejo E, et al. Tagging-SNP haplotype analysis of the secretory PLA2IIa gene PLA2G2A shows strong association with serum levels of sPLA2IIa: results from the UDACS study. Hum Mol Genet. 2006;15:355–61.

[35] Suhre K, Arnold M, Bhagwat AM, Cotton RJ, Engelke R, Raffler J, et al. Connecting genetic risk to disease end points through the human blood plasma proteome. Nat Commun. 2017;8:14357.

[36] Carayol J, Chabert C, Di Cara A, Armenise C, Lefebvre G, Langin D, et al. Protein quantitative trait locus study in obesity during weight-loss identifies a leptin regulator. Nat Commun. 2017;8:2084.

[37] Khan K, Zafar S, Hafeez A, Badshah Y, Shahid K, Mahmood Ashraf N, et al. PRKCE non-coding variants influence on transcription as well as translation of its gene. RNA Biol. 2022;19:1115–29.

[38] Hurt-Camejo E, Camejo G, Peilot H, Oorni K, Kovanen P. Phospholipase A(2) in vascular disease. Circ Res. 2001;89:298–304.

[39] Smith JW, Barlas RS, Mamas MA, Boekholdt SM, Mallat Z, Luben RN, et al. Association between serum secretory phospholipase A2 and risk of ischaemic stroke. Eur J Neurol. 2021;28:3650–5.

[40] Talvinen KA, Kemppainen EA, Nevalainen TJ. Expression of group II phospholipase A2 in the liver in acute pancreatitis. Scand J Gastroenterol. 2001;36:1217–21.

[41] Cai Y, Yu X, Hu S, Yu J. A brief review on the mechanisms of miRNA regulation. Genomics Proteomics Bioinformatics. 2009;7:147–54.

[42] Ramirez-Bello J, Jimenez-Morales M. [Functional implications of single nucleotide polymorphisms (SNPs) in protein-coding and non-coding RNA genes in multifactorial diseases]. Gac Med Mex. 2017;153:238–50.

[43] Consortium GT. The Genotype-Tissue Expression (GTEx) project. Nat Genet. 2013;45:580–5.

[44] Wang K, Li M, Hakonarson H. ANNOVAR: functional annotation of genetic variants from high-throughput sequencing data. Nucleic Acids Res. 2010;38:e164.

